# Unraveling the transcriptional networks that drive oligodendrocyte fate specification in Sonic hedgehog-responsive neocortical progenitors

**DOI:** 10.1101/2020.10.18.344515

**Authors:** Caitlin C. Winkler, Luuli N. Tran, Ellyn P. Milan, Fernando García-Moreno, Santos J. Franco

## Abstract

In the developing nervous system, progenitors first generate neurons before making astrocytes and oligodendrocytes. We previously showed that increased Sonic hedgehog (Shh) signaling in dorsal forebrain progenitors is important for their production of oligodendrocytes as neurogenesis winds down. Here, we analyzed single-cell RNA sequencing datasets to better understand how Shh controls this neuron-to-oligodendrocyte switch in the neocortex. We first identified Shh-responding progenitors using a dataset in which Shh was overexpressed in the mouse dorsal forebrain. Pseudotime trajectory inferences revealed a subpopulation committed to the oligodendrocyte precursor cell (OPC) lineage. Genes upregulated along this lineage defined a pre-OPC state, as cells transitioned from progenitors to OPCs. Using several datasets from wild-type mouse and human embryos at different ages, we confirmed a pre-OPC state preceding OPC emergence during normal development. Finally, we show that pre-OPCs are enriched for a gene regulatory network involving the transcription factor Ascl1. Genetic lineage-tracing demonstrated Ascl1^+^ dorsal progenitors primarily make oligodendrocytes. We propose a model in which Shh shifts the balance between opposing transcriptional networks toward an Ascl1 lineage, thereby facilitating the switch between neurogenesis and oligodendrogenesis.

## INTRODUCTION

Proper assembly of neural circuits in the developing brain requires spatially and temporally coordinated production of many different neuronal and glial cell types. In the developing dorsal forebrain, neural progenitor cells produce excitatory neurons, olfactory bulb interneurons, astrocytes and oligodendrocytes. Both internal and external factors are important for setting up this diversity, by translating spatial and temporal signaling cues into transcriptional outputs that determine cellular identity and function. A major goal in the field is to understand the molecular mechanisms that specify these different cell fates in the proper numbers and with appropriate timing.

The morphogen signaling molecule, Sonic hedgehog (Shh), is important for specifying oligodendrocyte fates in many different regions of the CNS. Studies from our lab previously showed that Shh signaling is both necessary and sufficient for the generation of neocortical oligodendrocytes from dorsal forebrain progenitors (Winkler et al., 2018). We proposed a model for how the timing and location of Shh signals affect the neuron-glia switch during neocortical development. In the early dorsal forebrain, Shh signaling is maintained at low levels to allow normal dorsal-ventral patterning. During this time, dorsal progenitors undergo neurogenesis to make neocortical excitatory neurons. At later stages, Shh ligand reaches the dorsal forebrain via infiltrating interneurons and circulating cerebral spinal fluid (Winkler et al., 2018). We showed that this increase in dorsal Shh is critical for instructing progenitors to begin oligodendrogenesis (Winkler et al., 2018) and to produce the full complement of oligodendrocyte lineage cells for the mature brain (Winkler and Franco, 2019). However, the intrinsic pathways that specify an oligodendrocyte fate downstream of Shh remain largely unknown.

In this study, we analyzed several publicly available single-cell RNA sequencing (scRNA-seq) datasets to investigate how Shh signaling influences progenitor cell fate decisions in the developing dorsal forebrain. We first analyzed data from a recent study that overexpressed Shh ligand during mid-corticogenesis in mouse embryos (Zhang et al., 2020). We identified the progenitors whose transcriptomes were enriched for components of the Shh pathway. We uncovered further diversity within these Shh-responding progenitors, with 2 main lineage trajectories predicted to generate either oligodendrocyte precursor cells (OPCs) or olfactory bulb interneurons (OB-INs). Within the OPC lineage, we identified a set of genes that defines a pre-OPC state between radial glial cells and OPCs. Using this gene set, we identified pre-OPCs in several other scRNA-seq datasets collected from unmanipulated wild-type mouse embryos at different stages of corticogenesis. We also identified a pre-OPC progenitor state in data from human fetal neocortex and found that human pre-OPCs share some common gene expression profiles with outer radial glial cells. Finally, we identified the transcriptional regulator, Ascl1, as a central component of a gene regulatory network enriched in pre-OPCs. We confirmed that *Ascl1* mRNA and protein are expressed in a subset of dorsal forebrain progenitors prior to OPC production, and demonstrate that Ascl1^+^ progenitors primarily generate oligodendrocytes at late embryonic ages. These data support a model of how increased Shh levels during late corticogenesis are translated by dorsal progenitors into a switch in cell fates between neurons and oligodendrocytes.

## RESULTS

### Identification of Shh-responding progenitor cells

To gain insights into how Shh signaling promotes oligodendrogenesis in the dorsal forebrain, we analyzed published scRNA-seq data in which Shh was ectopically expressed in the cortical ventricular zone (VZ) via *in utero* electroporation (IUE) at E13.5 (Zhang et al., 2020). Samples were collected at E16.5 for scRNA-sequencing analysis (Fig. 1A). This gave us a system to study the molecular pathways downstream of Shh signaling in dorsal forebrain neural progenitors. We first identified major cell types based on gene expression (Fig. 1, Table S1). As expected, most cells were excitatory projection neurons at various stages of development (Fig. 1A-B). Progenitor cells, which included radial glial cells (RGCs), intermediate progenitor cells (IPCs), and oligodendrocyte precursor cells (OPCs), represented 4 clusters (Fig. 1A-C). We also identified 2 distinct inhibitory interneuron (IN) clusters (Fig. 1A, C), one of which comprised INs generated in the ventral forebrain that had migrated into the neocortex. The other IN cluster was classified as precursors for olfactory bulb interneurons (OB-INs), based on differential expression of OB-IN markers.

**Fig. 1.**
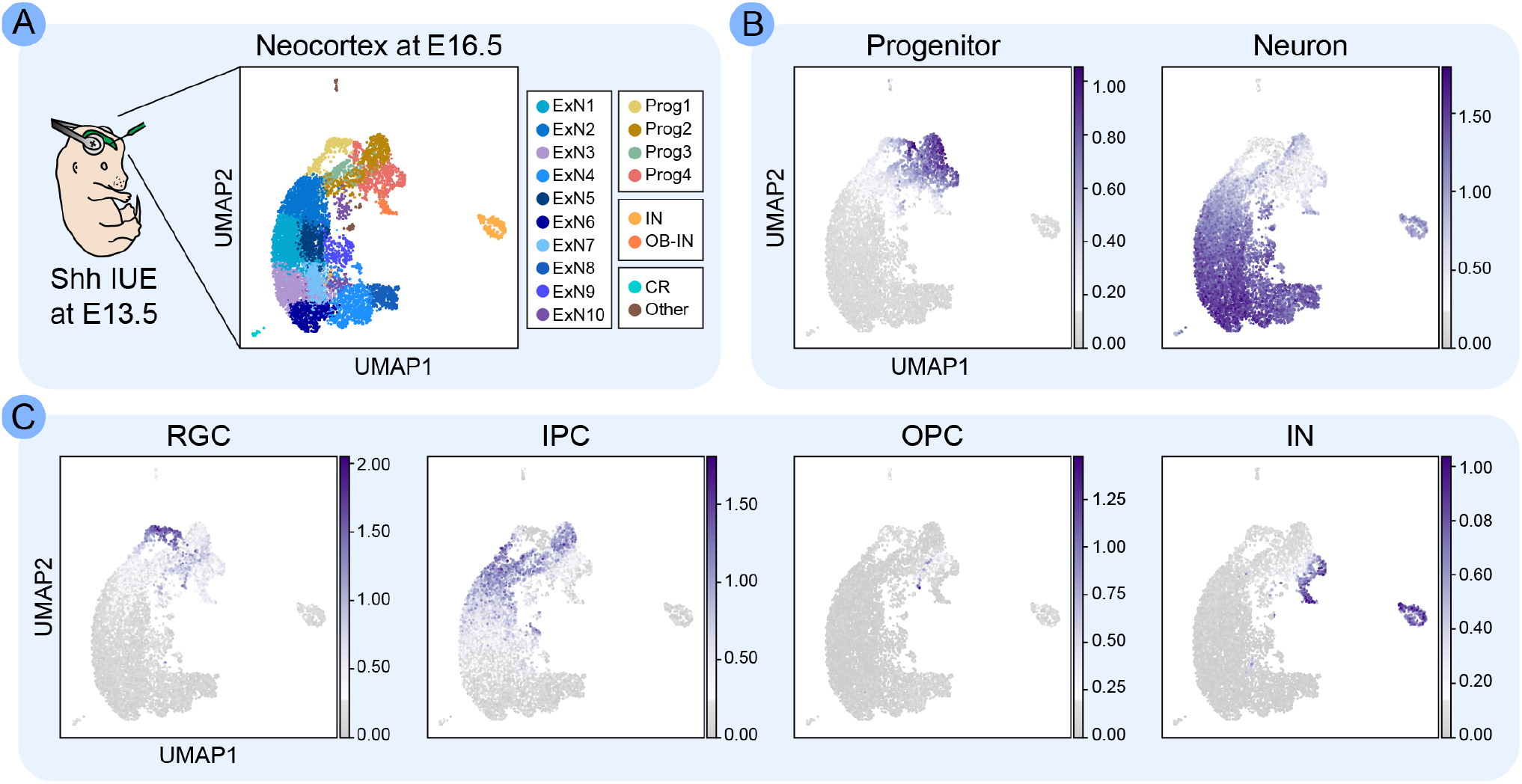
Identification of Progenitor Cells. (A) We analyzed data generated by Zhang et al., in which they ectopically expressed Shh in the cortical VZ by electroporating progenitors with *pCAG-ShhN-ires-GFP* plasmid at E13.5 (Shh IUE) and performed scRNA-seq at E16.5. UMAP visualization of resulting cell clusters identified by Scanpy and annotated by major cell type. “Other” denotes microglia and endothelial cells. (B) UMAP of cells colored by mean expression of Progenitor and Neuron marker gene sets. (C) UMAP of cells colored by mean expression of RGC, IPC, OPC and IN marker gene sets. See also Table S1 for marker genes.

We then re-clustered the progenitor cells for more detailed analysis. Because of their similarities to the progenitor clusters, we also included the immature OB-INs in our further analysis. Reclustering revealed 9 progenitor clusters, representing 5 different major cell types (Fig. 2A). The majority (56%) of cells were IPCs (*Eomes^+^*) at various stages of development toward excitatory neurons (Fig. 2A, B, D). RGCs (*Fabp7^+^*) constituted 15.5% of all progenitor cells (Fig. 2A, B, C). OB-INs (*Sp8^+^*) divided into two clusters, likely based on maturation state, and constituted 12.8% of all progenitors (Fig. 2A, B, F). OPCs (*Olig1^+^*) made up 2.8% of progenitors (Fig. 2A, E). The final progenitor cluster did not fall into any of these classifications (Fig. 2A-F). This cluster made up 12.9% of cells analyzed and was enriched for genes belonging to the Shh signaling pathway (Fig. 2G). Therefore, we named this cluster of cells Shh-responding progenitor cells, or ShhPCs.

**Fig. 2.**
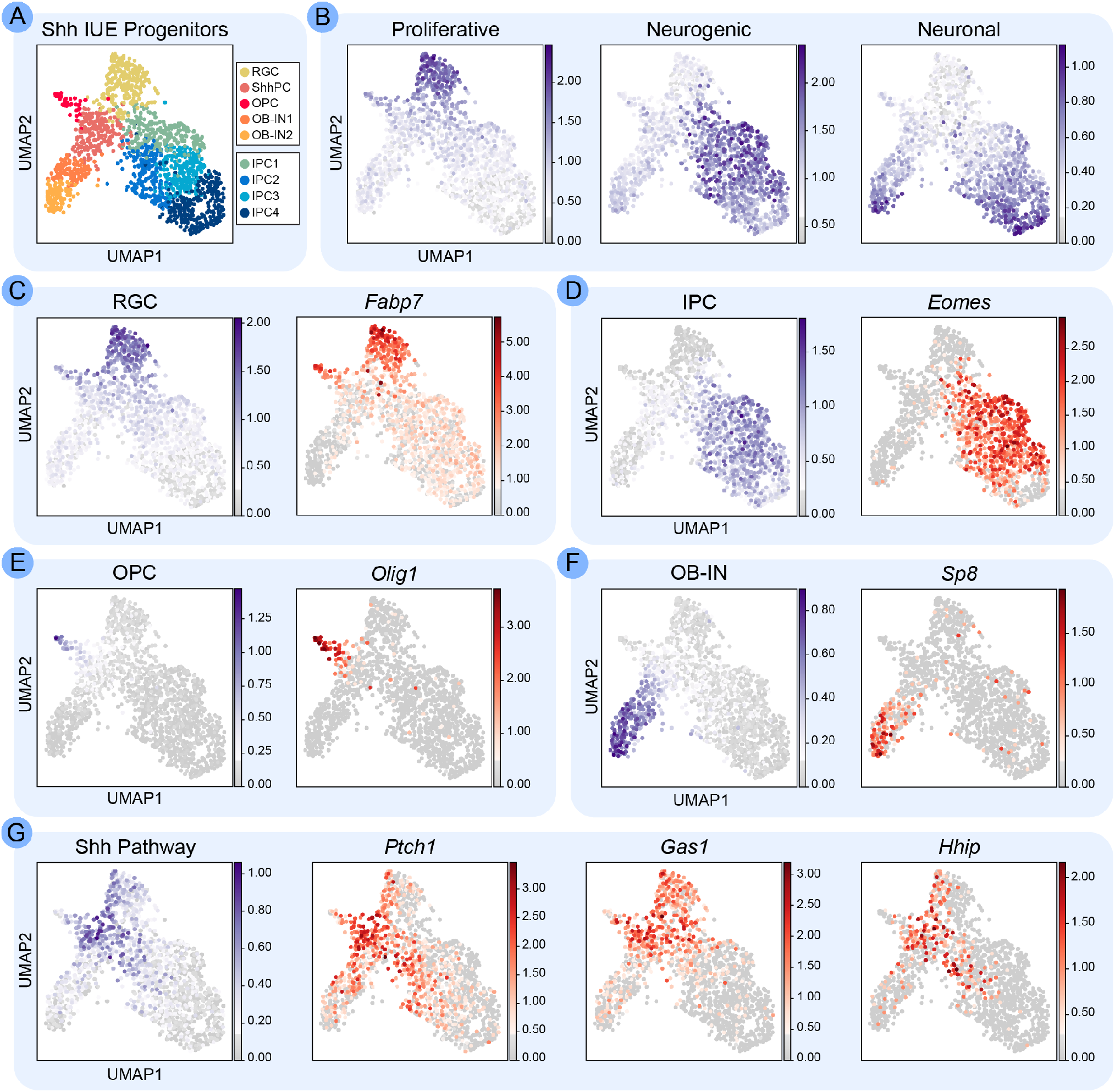
Clustering the Progenitor Cells. (A) Scanpy clustering was performed on all progenitor cells and presumptive OB-INs. Nine clusters were identified and annotated based on gene expression: RGC, radial glial cells; IPC1-4, intermediate progenitor cells; OB-IN1-2, olfactory bulb interneurons; OPC, oligodendrocyte precursor cells; ShhPC, Shh-responding progenitor cells. (B) UMAP of cells colored by mean expression of Proliferative, Neurogenic and Neuronal markers. (C-F) UMAPs of cells colored by mean expression of either groups of cell type markers (blue) or individual canonical marker genes (red) for each cell type. (G) UMAP of cells colored by mean expression of Shh Pathway markers (blue) and mean expression of individual genes (red). See also Table S1 for marker genes.

We hypothesized that the SthhPC cluster (Fig. 3A) was similar to the *Gsx2^+^* tri-potent progenitor cluster identified by Zhang et al. (Zhang et al., 2020). Indeed, we found that *Gsx2* mRNA was enriched in ShhPCs (Fig. 3B). We additionally identified several other differentially expressed genes enriched in the ShhPC cluster (Fig. 3B-C; Table S2), and others that were not only enriched but also fairly restricted to ShhPCs (Fig. 3D; Table S2). Of particular interest among the ShhPC differentially expressed genes was the transcription factor *Ascl1,* which is in the same transcriptional network as Gsx2 (Waclaw et al., 2009; Wang et al., 2009). Ascl1 plays an important role in oligodendrogenesis in the spinal cord and ventral forebrain (Nakatani et al., 2013; Parras et al., 2007; Sugimori et al., 2007) and is one of the first marker genes in the OPC lineage (Nakatani et al., 2013; Vue et al., 2014).

**Fig. 3.**
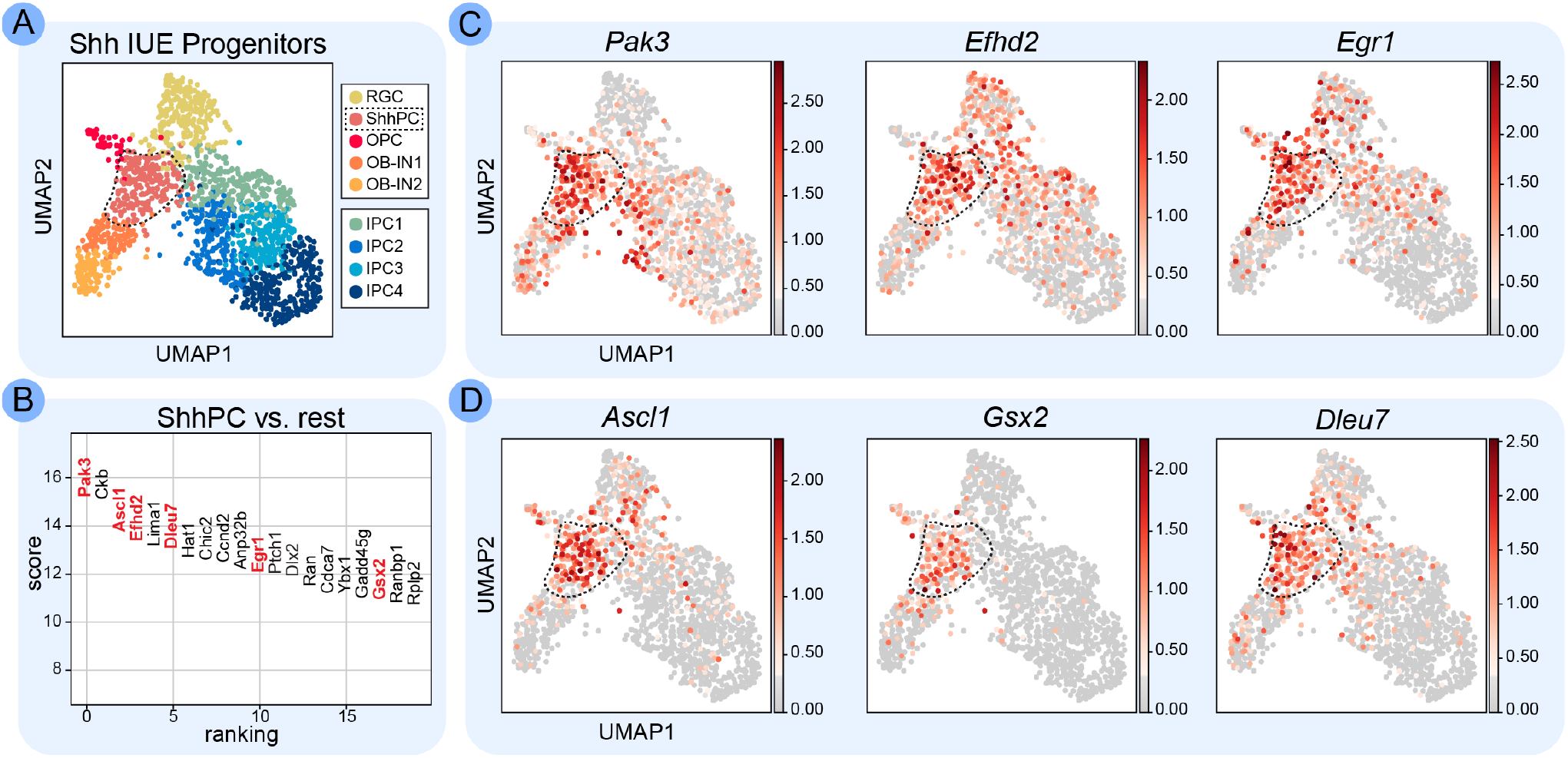
Differentially Expressed Genes in the Shh Progenitor Cluster. (A) UMAP of all progenitor cells and OB-INs, highlighting the ShhPC cluster. (B) Genes that are differentially expressed in the ShhPC cluster compared to all other clusters. Genes with corresponding UMAPs in (C) and (D) are shown in red. (C) UMAP of cells colored by mean expression of *Pak3, Efhd2* and *Egr1,* which are all enriched in the ShhPC cluster. (D) UMAP of cells colored by mean expression of *Ascl1, Gsx2* and *DIeu7,* which are all more restricted to the ShhPC cluster. See also Table S2 for ShhPC-enriched genes.

### Progenitor cluster relationships and lineage trajectories

To gain a deeper understanding of the relationships between progenitor clusters, we performed a heatmap and dendrogram analysis (Fig. 4A). We excluded cluster IPC1 as it was dominated by high expression of ribosome biogenesis genes, which confounded analysis. Based on hierarchical clustering predicted by the dendrogram, RGCs, ShhPCs, and OPCs were more closely related to each other than to OB-INs and IPCs (Fig. 4A). This is in line with ShhPCs and OPCs being more progenitor-like, whereas OB-INs and IPCs are more neuronal. Within the progenitor-like branch, ShhPCs and OPCs were predicted to be more closely related to each other than to RGCs, which is supported by shared expression of genes like *Ascl1.*

**Fig. 4.**
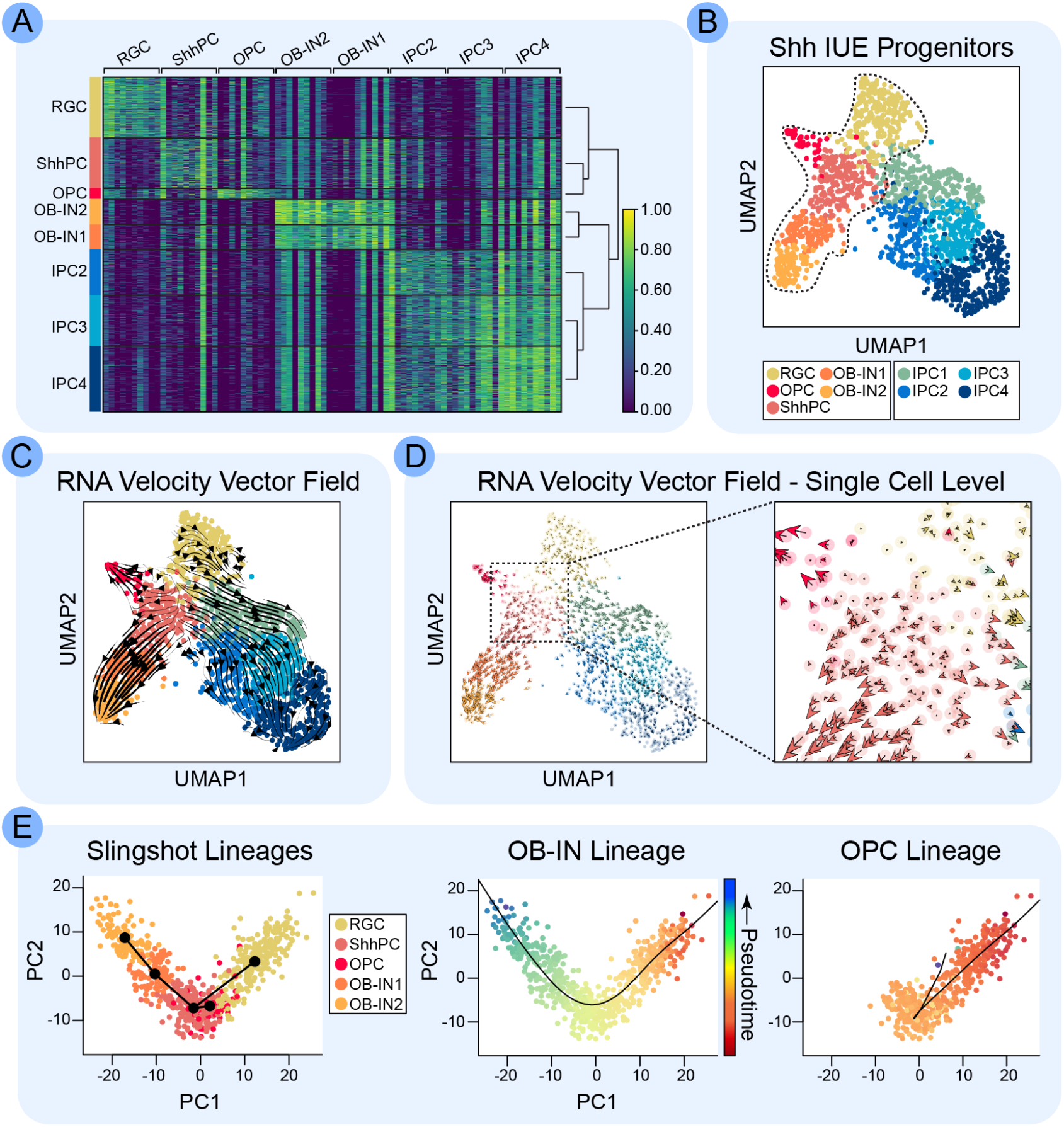
Lineage Analysis of OPCs and OB-INs. (A) Heatmap showing marker gene expression for the top genes of each cluster. Each column represents expression of one gene, and each row represents expression in one cell. Dendrogram shows hierarchical clustering. (B) UMAP of all progenitor cells and OB-INs, highlighting the arm that includes the RGC, ShhPC, OB-IN1, OB-IN2, and OPC clusters. (C) Velocities derived using the scVelo stochastic model, projected onto the UMAP. The main gene-averaged flow visualized by velocity streamlines shows a split between the IPC clusters and the OPC, OB-IN1 and OB-IN2 clusters, which are all derived from progenitors in the RGC and ShhPC clusters. (D) Fine-grained resolution of the velocity vector field in (C) shown at single-cell level Each arrow represents the direction and speed of movement of an individual cell. (E) Lineages of the clusters highlighted in (B), as predicted by Slingshot. Slingshot predicts that OPC and OB-IN lineages bifurcate at the ShhPC cluster. Cells are colored by cluster (left) or by pseudotime (middle and right). Note that the looping in the OPC lineage indicates that the endpoint would be coming into or out of the XY plane if presented in 3 dimensions.

To understand how the OPC lineage emerges during development, we focused on branches that included RGCs, ShhPCs, OPCs and OB-INs (Fig. 4B). We used RNA velocity (Bergen et al., 2020; La Manno et al., 2018) to predict lineage trajectories of cells in the different clusters (Fig. 4C). Projection of the velocities onto the progenitors indicated lineage progression from RGCs to IPCs, as expected (Fig. 4C). Additionally, RNA velocity analysis predicted cell-to-cell transitions from ShhPCs to both OPCs and OB-INs (Fig. 4C). Plotting the velocity vector fields at single-cell resolution allowed us to predict future states of individual cells (Fig. 4D). Cells in the more differentiated clusters of IPCs, OPCs and OB-INs displayed strong directional RNA velocity flow toward the ends of their branches. Conversely, velocities of cells in the RGC and ShhPC clusters were smaller and less directional, reflecting their less committed state (Fig. 4D). Interestingly, some ShhPCs had RNA velocities oriented toward OB-INs whereas others were directed toward OPCs (Fig. 4D), suggesting that the two lineages diverge at the ShhPC stage. We therefore performed lineage trajectory analysis using Slingshot (Street et al., 2018) on the RGC, ShhPC, OPC, and OB-IN clusters (Fig. 4B). Slingshot predicted two distinct lineages emerging from the ShhPC cluster (Fig. 4E). Both lineages started at RGCs and transitioned through the ShhPC state, at which point they diverged to either OPCs or OB-INs (Fig. 4E). Aligning predicted Slingshot lineages along pseudotime indicated that ShhPCs were between RGCs and the more differentiated OB-IN or OPC clusters (Fig. 4E), reinforcing the idea that ShhPCs are progenitors that are committing to these two lineages in response to Shh signaling.

We next employed tradeSeq (Van den Berge et al., 2020) to identify differentially expressed genes between the OPC and OB-IN lineages over pseudotime (Fig. 5, Table S3). We focused our analysis between tradeSeq knots 2 and 5 (see Methods), as this incorporated ShhPCs and the predicted divergence of the OPC and OB-IN lineages (Fig. 5A). Estimated smoother curves for the known OPC-specific gene, *Olig1*, showed upregulated and stabilized expression in the OPC lineage over pseudotime (Fig. 5B). Conversely, smoother curves for the OB-IN specification gene, *Sp8*, indicated sustained expression in the OB-IN lineage (Fig. 5B). Beyond these well-characterized examples of OPC- or OB-IN-specific markers, we identified several genes expressed in ShhPCs that either stayed on or were further upregulated in the OPC lineage, but not in the OB-IN lineage (Fig. 5C). Interestingly, *Ascl1* and *Gsx2* were maintained in newly formed OPCs but downregulated in newly formed OB-INs (Fig. 5C), implicating these transcription factors specifically in the OPC lineage. In contrast, several other genes were upregulated or maintained in the OB-IN lineage and downregulated in the OPC lineage (Fig. 5D). These included genes important for OB-IN fate determination, such as the transcription factors *Sp9, Pax6* and *Dlx1* (Fig. 5D). Together, these lineage analyses allowed us to identify the transcriptional changes conferred by Shh signaling that prime a subset of dorsal forebrain progenitors to acquire OPC or OB-IN fates.

**Fig. 5.**
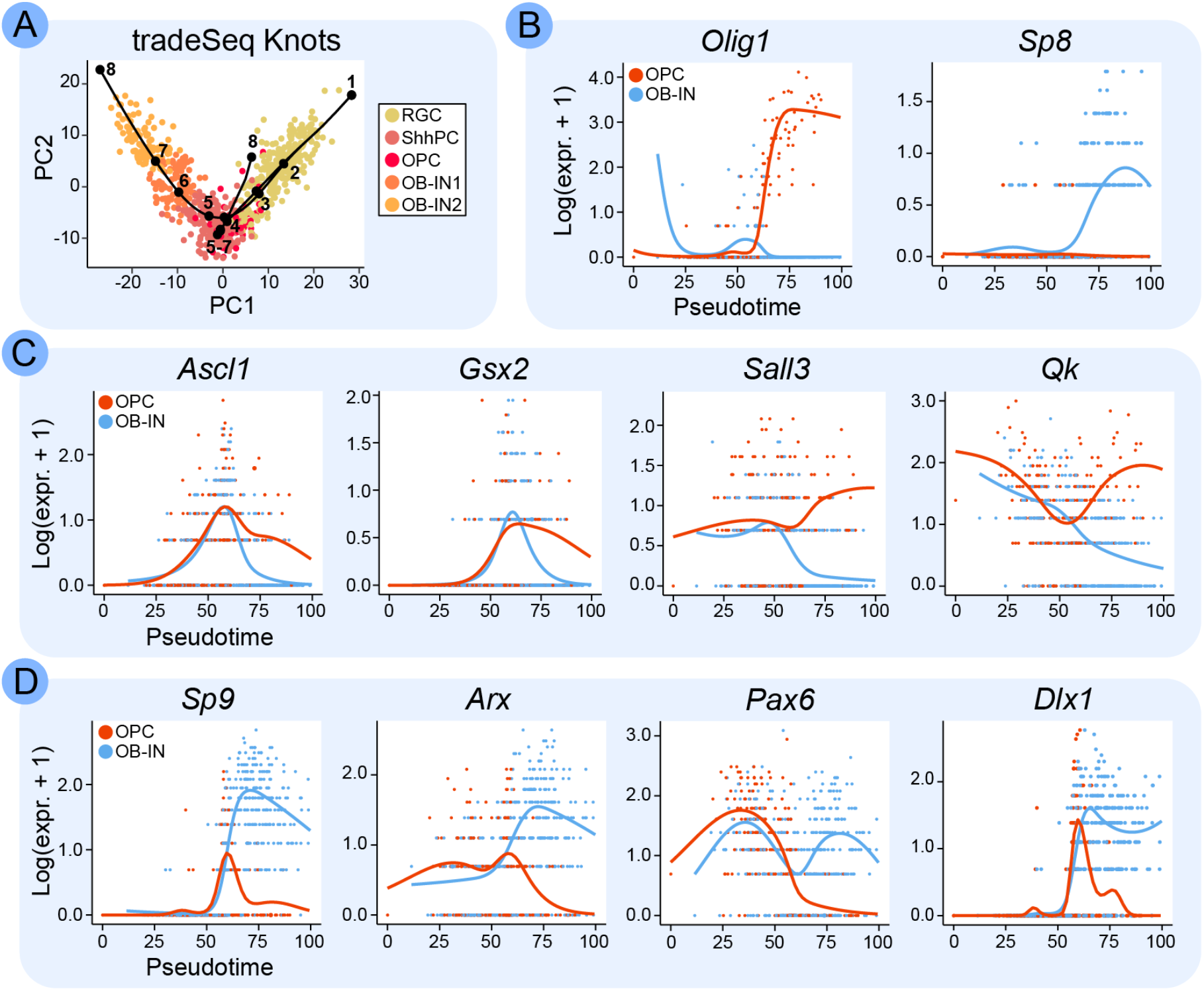
Differentially Expressed Genes Between OPC vs. OB-IN Lineages. (A) PCA plot for lineage trajectories predicted by Slingshot, illustrating the 8 knots used for tradeSeq analysis. tradeSeq was used to identify genes differentially expressed between the OPC and OB-IN lineages between knots 2 and 5. Cells are colored by cluster. (B) Estimated smoothers for *Olig1* and *Sp8,* as predicted by tradeSeq. (C) Estimated smoothers for example OPC lineage-specific genes predicted by tradeSeq. *Ascl1, Gsx2, Sall3,* and *Qk* all have predicted expression in ShhPCs, and continued expression in the OPC lineage over pseudotime. (D) Estimated smoothers for example OB-IN lineage-specific genes predicted by tradeSeq. *Sp9, Arx, Pax6,* and *Dlx1* all have predicted expression in ShhPCs, and continued expression in the OB-IN lineage. See also Table S3 for tradeSeq results.

Given the clear split in gene expression over pseudotime between the OPC and OB-IN lineages, we further subclustered the ShhPCs to identify any subpopulations (Fig. 6A). We found 3 distinct ShhPC subclusters: ShhPC1, ShhPC2, and ShhPC3. Similar to cluster IPC1, ShhPC3 was dominated by high expression of genes involved in ribosome biogenesis. Therefore, we focused on ShhPC1 and ShhPC2. Enrichment analysis of ShhPC1 found overrepresentation of genes with gene ontology (GO) terms associated with gliogenesis, glial fate commitment, and oligodendrocyte differentiation (Fig. 6B, Tables S4 and S5). Alternatively, ShhPC2 had over-representation of genes with GO terms associated with neuron fate commitment and interneuron differentiation (Fig. 6B, Tables S6 and S7). Further examination of specific gene expression reflects this heterogeneity (Fig. 6C); ShhPC1 had higher expression in genes associated with Notch signaling (*Hesi, Notchi),* Shh signaling (*Ptchi, Hhip, Gasi),* and OPC function (*Olig2, Pdgfra*) compared to ShhPC2, which showed higher expression in genes associated with the OB-IN lineage (*Dlx1, Dlx2, Arx, Sp9).* Interestingly, ShhPC1 shared some weak gene expression with RGCs (*Ptprz1*, *Hes1*, *Notch1*), while ShhPC2 shared *Gadd45g* expression with IPCs, perhaps indicating that ShhPC1 cells retain more RGC-like characteristics and ShhPC2 cells are more IPC-like (Fig. 6C).

**Fig. 6.**
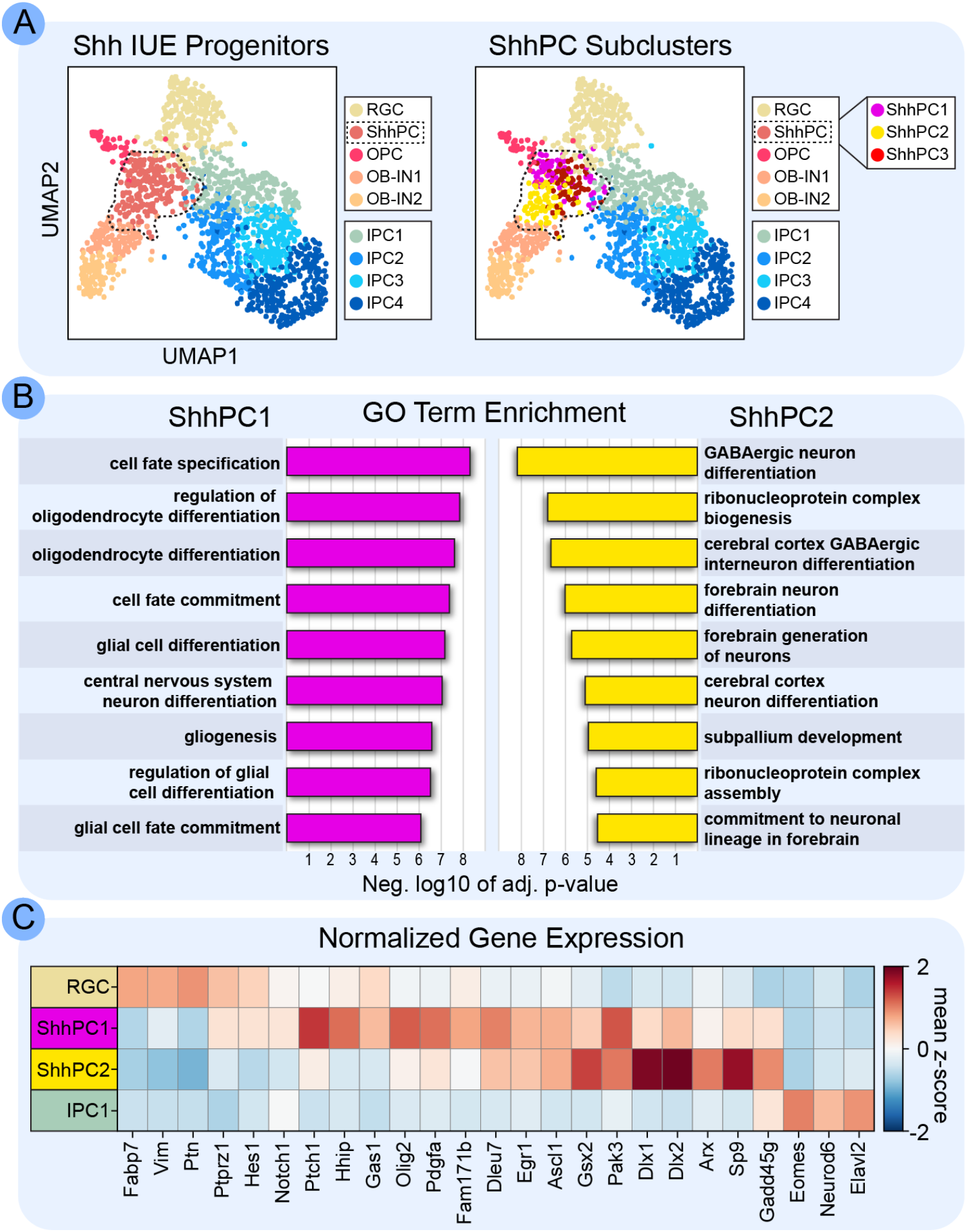
Subclustering the Shh Progenitor Cluster. (A) UMAP of all progenitor cells and OB-INs, highlighting the ShhPC cluster (left). The ShhPC cluster subclustered into three distinct subclusters (right): ShhPC1, ShhPC2, and ShhPC3. (B) Functional annotation of differentially expressed genes identified by Scanpy show top Gene Ontology (GO) term biological processes associated with ShhPC1 and ShhPC2. (C) Heatmap showing expression of specific genes of interest between ShhPC1 and ShhPC2, using RGC and IPC1 as control comparisons. See also Tables S4-7 for enriched genes and GO term results.

Together, our analyses indicate that increased Shh signaling can instruct dorsal forebrain RGCs towards a Shh-responsive progenitor cell fate, and that these ShhPCs can generate OPC and OB-IN lineages. We identified pre-OPCs as a subpopulation of ShhPCs that appear to be committed to the oligodendrocyte lineage and precede OPCs based on gene expression: pre-OPCs express genes shared between all ShhPCs in addition to some that are more restricted. Thus, we were able to generate a combinatorial gene expression signature for pre-OPCs.

### Identification of pre-OPCs in wild-type brains

Because these data involved ectopic expression of Shh, we wondered whether the pre-OPC signature could be found in normally developing brains. Therefore, we analyzed scRNA-seq data collected from the neocortices of wild-type mice at different embryonic ages (Fig. 7). We first analyzed a dataset from E18.0 dorsal forebrains (La Manno et al., 2020), since this is a stage when dorsally-derived cells in the OPC lineage are abundant (Winkler et al., 2018). We clustered all cells based on gene expression to identify progenitors (Fig. 7A), then focused our analysis on just the progenitor clusters. Classification of progenitor clusters based on gene expression profiles identified RGCs, IPCs, INs, astrocytes and OPCs (Fig. 7A-B). Overlaying our putative pre-OPC signature clearly identified clusters with high expression of this gene set (Fig. 7B). Cells with a strong pre-OPC gene expression signature were adjacent to cells in the RGC and OPC clusters that exhibited weaker expression of pre-OPC genes (Fig. 7B). Individual genes from the pre-OPC signature were also enriched, including *Ascl1, Gsx2, Egfr* and *Sall3* (Fig. 7C). These data indicate that the pre-OPC transcriptional profile is present in a subset of progenitors in the E18.0 wild-type dorsal forebrain.

**Fig. 7.**
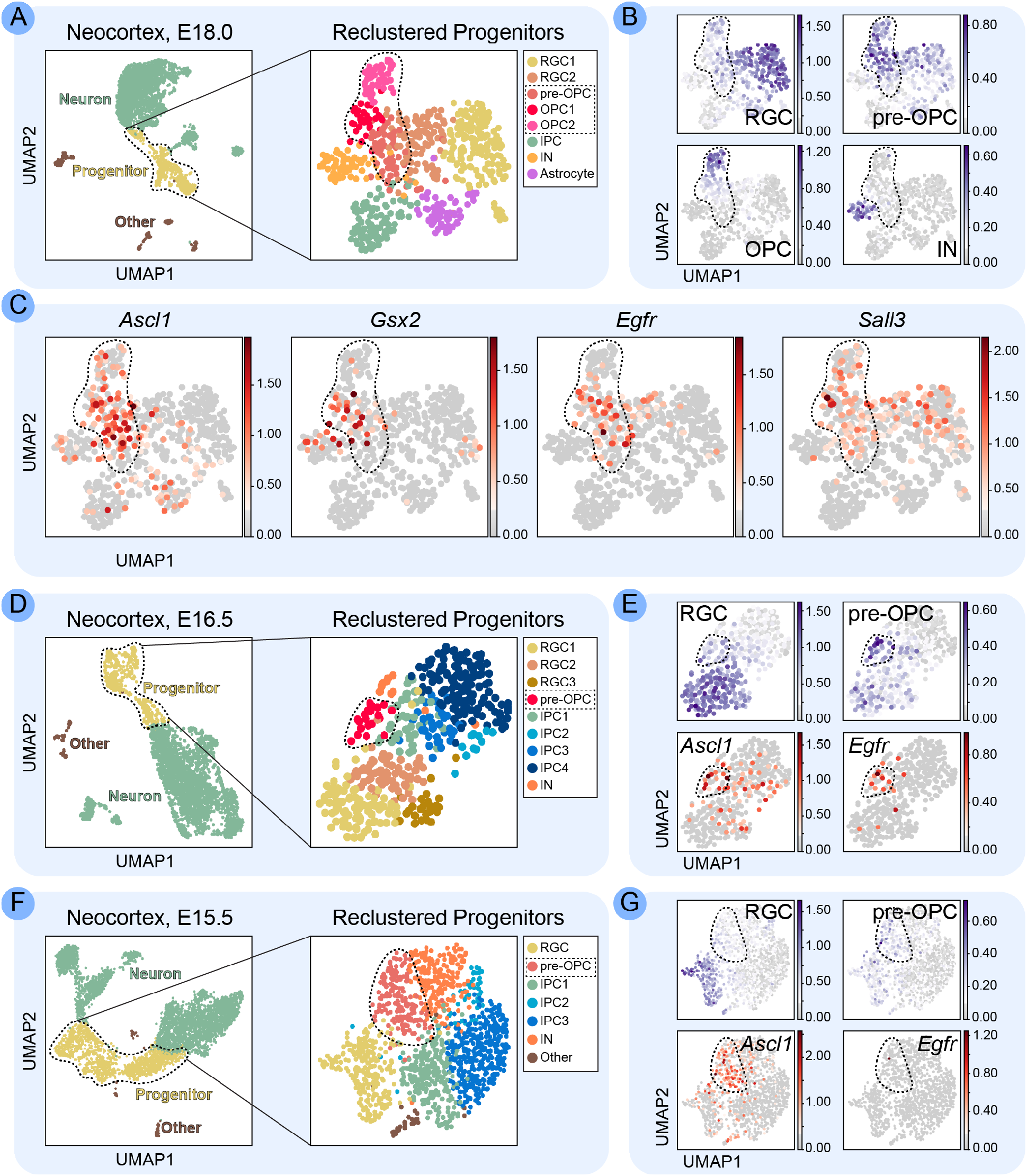
Pre-OPC Progenitors in Normal Development. (A) Scatterplot of cells after principal-component analysis and UMAP visualization of E18.0 wildtype (WT) neocortex. Left panel: all cells collected for scRNA-seq analysis. Right panel: new principal-component analysis and UMAP visualization of just the progenitor clusters. Eight progenitor clusters were identified and annotated based on gene expression: radial glial cells (RGC1-2), oligodendrocyte precursor cells (pre-OPC, OPC1-2), intermediate progenitor cells (IPC), interneurons (IN) and Astrocytes. Cells are colored according to Scanpy clustering. (B) UMAP of E18.0 progenitor cells colored by mean expression of RGC, pre-OPC, OPC and IN marker gene sets. (C) UMAP of E18.0 progenitor cells colored by mean expression of specific pre-OPC genes identified in the E16.5 Shh-IUE dataset. (D) Same as in (A), except with E16.5 WT neocortex data. (E) UMAP of E16.5 progenitor cells colored by mean expression of RGC and pre-OPC marker gene sets (blue), and by mean expression of *Ascl1* and *Egfr* (red). (F) Same as in (A) except with E15.5 WT neocortex data. (G) Same as in (E) except with E15.5 progenitor cells. For all UMAPs, “Other” denotes microglia and endothelial cells.

We next analyzed scRNA-seq data from E16.5 (Fig. 7D-E) (La Manno et al., 2020) and E15.5 (Fig. 7F-G) (Yuzwa et al., 2017). Very few OPCs are present at these timepoints, but a cluster of progenitor cells at each age exhibited robust expression of genes in the pre-OPC signature (Fig. 7E, G). At both ages, expression of *Ascl1* was enriched in the putative pre-OPC clusters (Fig. 7E, G). *Egfr* was previously identified as being enriched early in OPC development (Falcão et al., 2018; Huang et al., 2020; La Manno et al., 2020; Marisca et al., 2020; Marques et al., 2016). We found that *Egfr* expression was enriched in pre-OPCs at E18.5 (Fig. 7C) and E16.5 (Fig. 7E) but was not yet strongly expressed at E15.5 (Fig. 7G). Thus, *Ascl1;Egfr* progenitors at E15.5 could be ShhPCs that have not yet committed to the pre-OPC lineage, which would be consistent with studies from our lab that dorsal progenitors start to give rise to OPCs between E15.5-E17.5 (Winkler et al., 2018).

### Validation of Ascl1 as a pre-OPC marker in the embryonic dorsal forebrain

Previous studies demonstrated important roles for *Ascl1* in OPC development in the spinal cord (Battiste et al., 2007; Kelenis et al., 2018; Sugimori et al., 2007; Sugimori et al., 2008; Vue et al., 2014), embryonic ventral forebrain (Parras et al., 2007; Yung et al., 2002) and postnatal telencephalon (Nakatani et al., 2013; Parras et al., 2004). These studies indicate that *Ascl1* is already expressed early in OPC development (Battiste et al., 2007; Parras et al., 2007; Sugimori et al., 2007). To validate our scRNA-seq analysis, we performed multiplex mRNA *in situ* hybridization to detect *Ascl1* mRNA and the dorsal progenitor marker, *Emx1* (Fig. 8A-B). At E15.5 *Emx1* was highly expressed in the dorsal progenitor zones, while *Ascl1* expression was strongest in ventral progenitors (Fig. 8A). However, we could readily detect low levels of *Ascl1* in the *Emx1* VZ, where dorsal progenitors reside (Fig. 8A). By E16.5, *Ascl1* expression was even more evident in the *Emx1* dorsal VZ, although it was still weaker than in ventral regions (Fig. 8B). We next performed immunohistochemistry for Ascl1 protein in E16.5 *Emx1-Cre;NZG* mice (Winkler et al., 2018), in which dorsal RGCs and their progeny express nuclear β-gal (Fig. 8C). We detected Ascl1;β-gal double-positive cells in the ventricular and subventricular zones, further confirming Ascl1 expression in dorsal progenitors. Ascl1 expression was weak in the VZ and stronger in more basal cells (Fig. 8C), suggesting expression is initiated in RGCs. Interestingly, apical cells that were weakly positive for Ascl1 were usually negative for Olig2, whereas the higher-expressing basal Ascl1^+^ cells often co-expressed Olig2 (Fig. 8C). This pattern is consistent with Ascl1 expression being initiated in RGCs before Olig2 expression starts, prior to acquisition of an OPC fate. We also identified a subset of basally-located cells strongly expressing Ascl1 but not Olig2 (Fig. 8C), which could represent Ascl1^+^ cells fated to make OB-INs instead of OPCs.

**Fig. 8.**
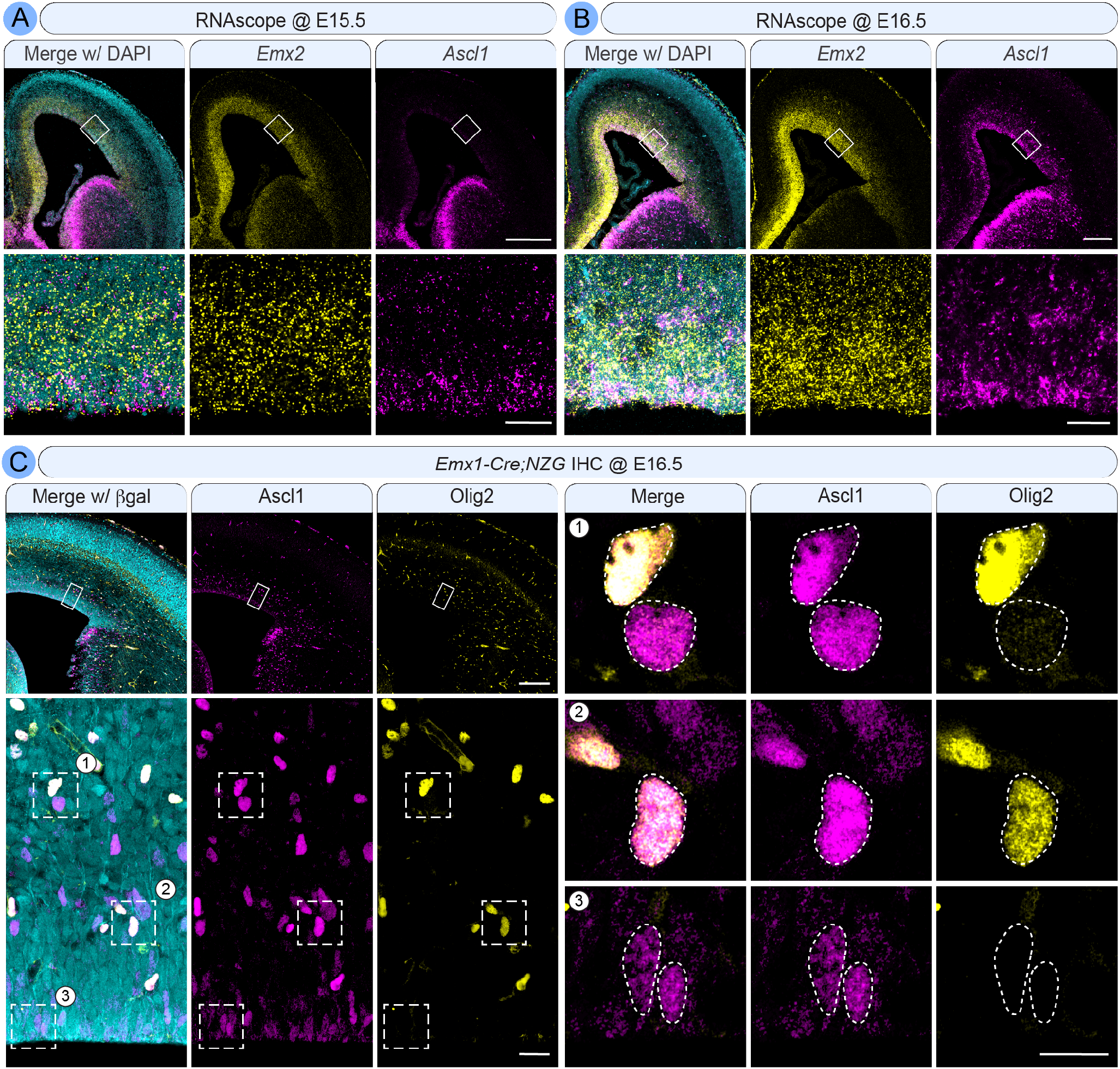
Expression of *Ascl1* mRNA and Ascl1 protein in *Emx1^+^* dorsal progenitors. (A-B) RNAscope® assay to analyze mRNA levels of *Emx2* (yellow) and *Ascl1* (magenta) in the neocortex at E15.5 (A) and E16.5 (B), counterstained with DAPI (cyan) for nuclei. Top rows: overviews showing *Emx1* expression is highest in dorsal progenitor domains, whereas *Ascl1* is enriched in ventral regions. Bottom rows: higher magnification of the boxed areas showing *Ascl1* is also expressed in the *Emx1^+^* dorsal progenitor domain, and appears to increase from E15.5 (A) to E16.5 (B). Scale bars: 200 μm overview (top rows), 20 μm zoom (bottom rows). (C) Immunohistochemistry to analyze Ascl1 (magenta) and Olig2 (yellow) protein at E16.5 in *Emx1-Cre;NZG* reporter mice. β-gal (cyan) labels all cells from the Emx1+ dorsal progenitor lineage. Upper left panels: an overview of the imaged region. Bottom left panels: higher magnification of the boxed regions in the overviews, to identify β-gal/Ascl1 double-positive cells (numbered boxes). Panels 1-3 on the right are zoomed-in images of the corresponding numbered boxes on the bottom left, showing: 1) Two Ascl1^+^ nuclei in the upper subventricular zone, one co-expressing high levels of Olig2 and the other Olig2-negative; 2) Two Ascl1^+^ nuclei in the lower subventricular zone, both expressing moderate levels of Olig2; 3) Two nuclei in the ventricular zone weakly expressing Ascl1, both Olig2-negative. Scale bars: 200 μm overview (top left row), 20 μm zoom (bottom left row), 10 μm individual nuclei (right).

Lineage-tracing studies using *Ascl1-Cre* and *Ascl1-CreERT2* mice demonstrated that Ascl1^+^ progenitors give rise to many different cell types in the brain, including oligodendrocytes and OB-INs, but not neocortical excitatory neurons (Kelenis et al., 2018; Kim et al., 2008; Kim et al., 2011). However, these studies did not determine which cell types derived from Ascl1^+^ progenitors specifically in the dorsal forebrain. To test if dorsal forebrain Ascl1^+^ progenitors can generate neocortical oligodendrocytes, we lineage traced these progenitors using *in utero* electroporation of *Ascl1-CreERT2* embryos (Fig. 9). We crossed *Ascl1-CreERT2* mice (Kim et al., 2011) to the cre-reporter strain, *Ai9*(Madisen et al., 2010). We then used directional *in utero* electroporation of a *piggyBac* transposase-based plasmid (Garcia-Moreno et al., 2014) to achieve stable integration of a fluorescence reporter transgene (nuc.mT-Sapphire) specifically into dorsal RGCs at E15.5 (Fig. 9A). Immediately after *in utero* electroporation, we injected the pregnant dam with tamoxifen to label the Ascl1^+^ lineage with tdTomato (Fig. 9A). In this experiment, cells that are nuc.mT-Sapphire;tdTomato double-positive are specifically from the dorsal Ascl1^+^ lineage. At postnatal day P0, the majority of electroporated nuc.mT-Sapphire;tdTomato double-positive cells were located in the SVZ (Fig. 9B), although cells could be found throughout the neocortex. We could readily detect many nuc.mT-Sapphire;tdTomato double-positive that also expressed Olig2 (Fig. 9B, arrows), demonstrating that dorsal Ascl1^+^ progenitors can generate oligodendrocytes. We also identified a smaller number of nuc.mT-Sapphire;tdTomato double-positive cells that did not express Olig2 (Fig. 9B, arrowhead). The identity of these cells is not yet known, but they could be newborn OB-INs or possibly pre-OPCs that have not yet turned on Olig2 expression. Quantification of nuc.mT-Sapphire;tdTomato double-positive cells showed that 68.64%(± 3.48%, SEM) were Olig2^+^, indicating that oligodendrocytes are the primary cell type made by Ascl1^+^ dorsal progenitors at this age.

**Fig. 9.**
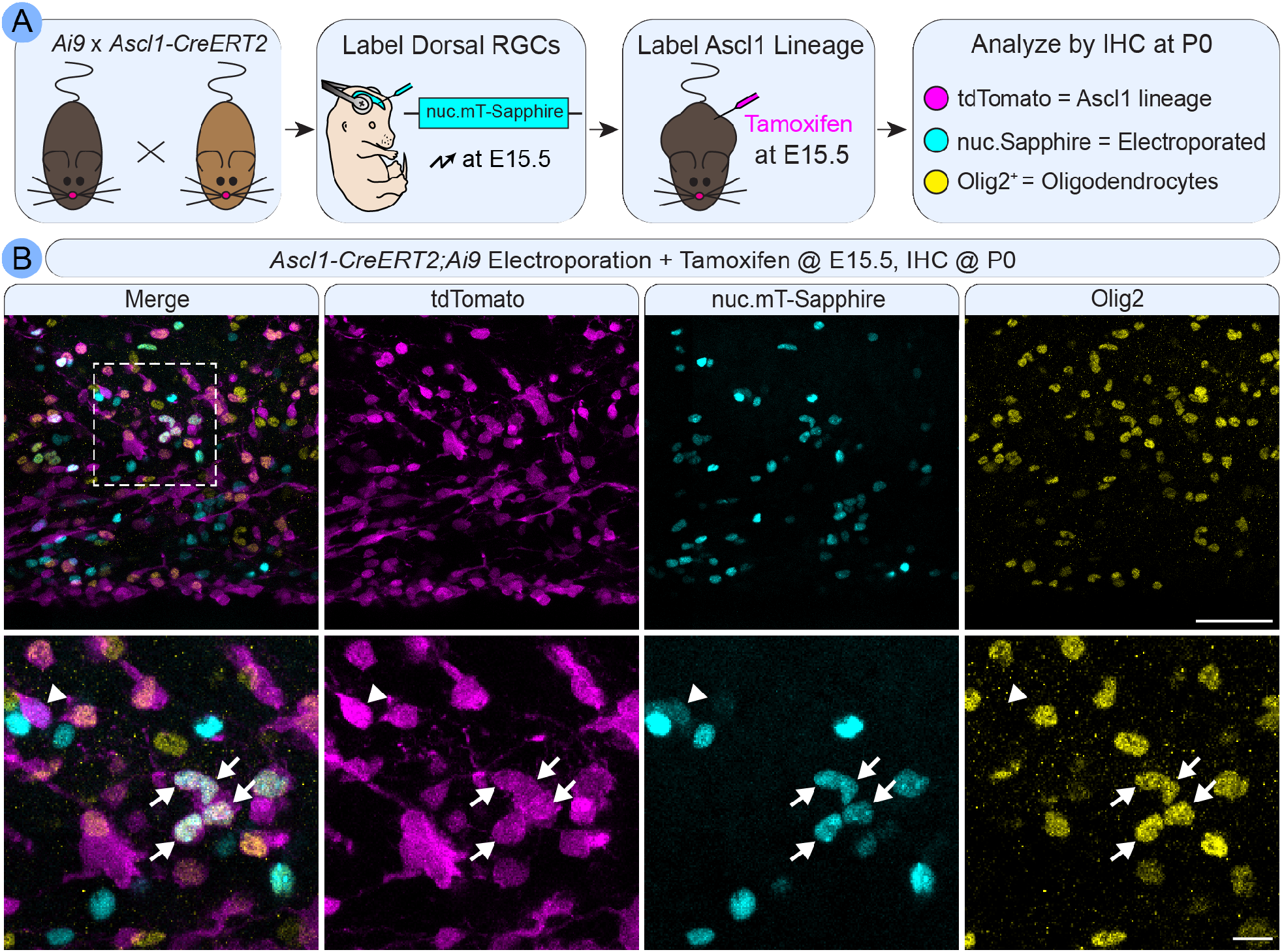
Dorsal Ascl1^+^ cells can make OPCs. (A) Schematic of the genetic lineage-tracing approach to characterize dorsally-derived Ascl1-lineage cells in the neocortex. *Ascl1^+/creERT2^* mice were crossed to *Ai9^fl/fl^* reporter mice. At E15.5, we used *in utero* electroporation to label dorsal RGCs and their offspring with nuc.mT-Sapphire. We then injected tamoxifen to permanently label all Ascl1^+^ progenitors and their offspring with tdTomato. Therefore, tdTomato/nuc.mT-Sapphire double-positive cells were from the dorsal Ascl1 lineage. Brains were collected at P0 and stained for Olig2 to label OPCs. (B) Immunohistochemical analysis of electroporated brains at P0. Top row, overview of the ventricular/subventricular zones showing tdTomato (magenta), nuc.mT-Sapphire (cyan) and Olig2 (yellow). Scale bar: 50 μm. Bottom row, higher magnification of cells from the boxed region. Arrows denote tdTomato/nuc.mT-Sapphire double-positive cells that are Olig2^+^, indicating OPCs derived from dorsal Ascl1^+^ RGCs. Arrowhead denotes a tdTomato/nuc.mT-Sapphire double-positive cell that is Olig2-negative. Scale bar: 10 μm.

### pre-OPCs in developing human brains

Finally, we wondered if the pre-OPC state identified in mice is conserved in the developing human neocortex. We analyzed data collected from human neocortices at gestational week 18 (Bhaduri et al., 2020), a time developmentally comparable to the end of neurogenesis and beginning of oligodendrogenesis in mice (E16.5-E18.5). Cells collected from various regions of the neocortex were batch corrected before analysis (Fig. 10A). Similar to the mouse datasets, the vast majority of human cells were excitatory neurons (Fig. 10B, Table S8). We identified 2 progenitor clusters, which were re-clustered together for further analysis (Fig. 10B). We found 14 distinct progenitor clusters, the majority of which were RGCs and IPCs (Fig. 10B). We also identified 3 distinct OPC clusters that separated based on maturational stages (Fig. 10B-C). All OPC clusters expressed pan-oligodendrocyte lineage markers *Olig1* and *OLIG2*, although half of the OPC1 cluster did not express *Olig1* or *OLIG2*(Fig. 10D). OPC3 appeared to be committed oligodendrocyte precursors (COPs) based on expression of genes like *BCAS1* and *SIRT2* (Fig. 10C, E). OPC2 was identified as true OPCs based on expression of genes like *SERPINE2* and *PDGFRA* (Fig. 10C, E). Some of these OPC markers were also weakly expressed in part of the OPC1 cluster (Fig. 10E). Because none of these known oligodendrocyte lineage markers fully covered the OPC1 cluster, we hypotHes1zed that this cluster might predominantly comprise pre-OPCs. Indeed, several pre-OPC genes identified in mice nicely incorporated the OPC1 cluster (Fig. 10F). Interestingly, none of the pre-OPC genes were exclusive to OPC1; rather, they either spanned the OPC1 and RGC clusters (*e.g., EGFR* and *RFX4*) or they spanned all OPC clusters and some RGC clusters (*e.g.*, *BCAN* and *GNGi2*) (Fig. 10F). Many of these pre-OPC genes were enriched specifically in RGC1 (Fig. 10B, F), which were identified as outer radial glial cells (oRGs). oRGs preferentially express *Ptprz1* and *HOPX* (Pollen et al., 2015), which are both expressed in the OPC1 cluster (Fig. 10G). *PTRPZ1* was enriched in the RGC1, OPC1, and OPC2 clusters, while *HOPX* was even further restricted to predominantly the RGC1 and OPC1 clusters (Fig. 10G). We also found *CD9* expression to be relatively restricted to the RGC1 and OPC1 clusters (Fig. 10G). oRGs also express genes associated more with astrocytes later in development, like *DIO2* (Pollen et al., 2015), which we found in both the RGC1 and astrocyte clusters, but not in the OPC clusters (Fig. 10G). Taken together, our analysis of the developing human neocortex identified a potential pre-OPC stage in which progenitors express low levels of some OPC markers and several pre-OPC markers identified in mice, indicating conservation of certain aspects of the pre-OPC state from mice to humans.

**Fig. 10.**
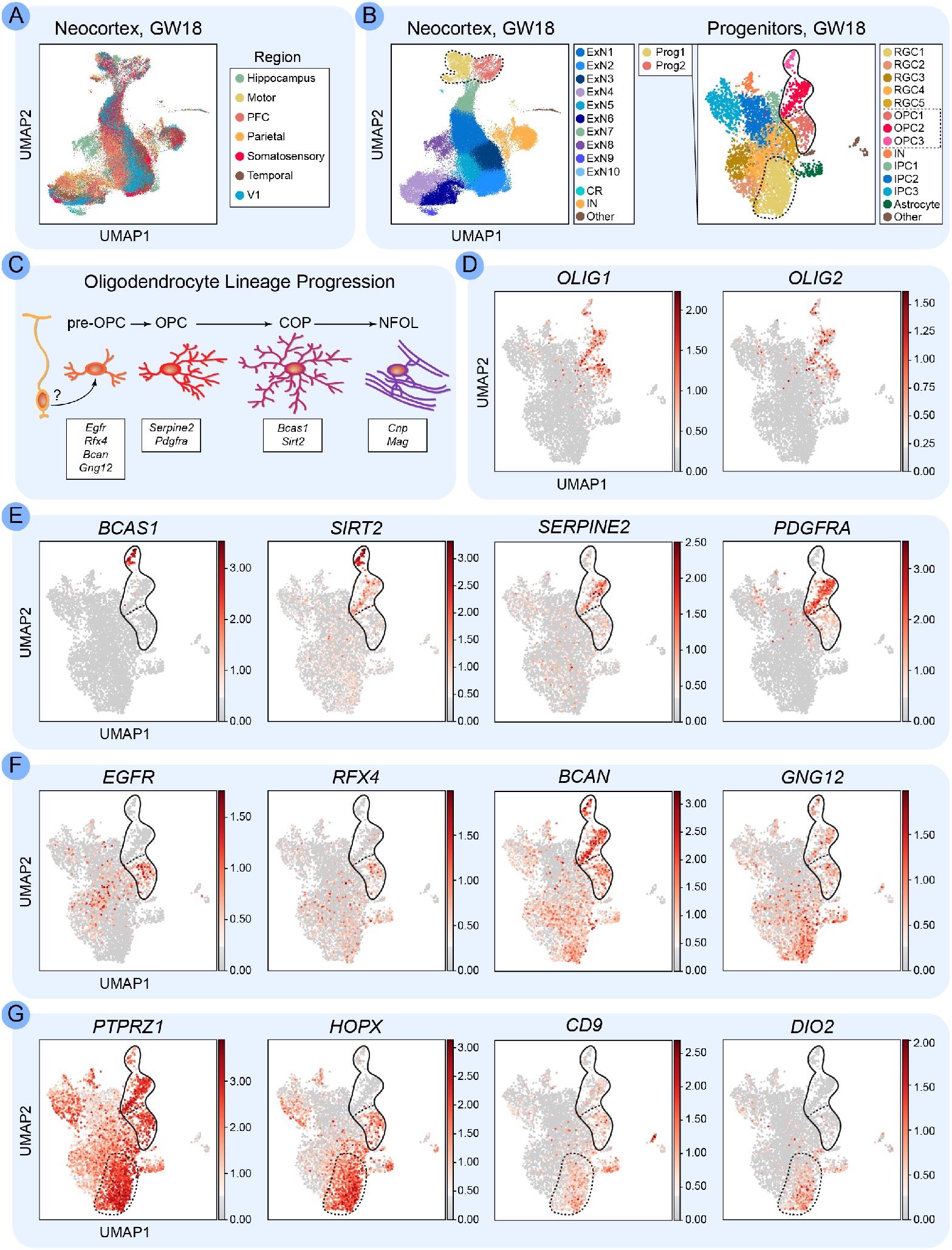
Identification of pre-OPCs in the human fetal neocortex. (A) Scatterplot of cells after principalcomponent analysis and UMAP visualization of human neocortical cells from gestational week 18. Several different neocortical regions were sampled and the data were batch corrected accordingly. (B) Left: UMAP visualization of cell clusters identified by Scanpy and annotated by major cell type. “Other” denotes microglia and endothelial cells. Right: resulting UMAP visualization after re-clustering the progenitor and OPC clusters. Fourteen clusters were identified and annotated based on gene expression: radial glial cells (RGC1-5); oligodendrocyte precursor cells (pre-OPC, OPC, COP); interneurons (IN); intermediate progenitor cells (IPC1-3); Astrocytes; Other (microglia and endothelial cells). (C) Schematic representing distinct stages of oligodendrocyte development. Neural progenitor cells give rise to pre-OPCs (characterized by *EGFR, RFX4, BCAN* and *GNG12* expression). pre-OPCs mature into OPCs (characterized by *SERPINE2* and *PDGFRA),* which produce committed precursors (COPs, characterized by *BCAS1* and *SIRT2*) that develop into newly formed oligodendrocytes (NFOL, characterized by *CNP* and *MAG).* (D-G) UMAPs of progenitor cells colored by mean expression of: (D) *OLIG1* and *OLIG2;* (E) COP and OPC markers; (F) pre-OPC markers; (G) pre-OPC/oRG-enriched gene *PTPRZ1* and oRG markers *HOPX, CD9,* and *DIO2.* Some oRG markers were exclusive to the RGC1 cluster (*DIO2*), whereas others were also expressed in the pre-OPC cluster (*PTPRZ1, HOPX, CD9).* See also Table S8 for marker genes.

## DISCUSSION

Neural progenitors in the mammalian dorsal forebrain generate diverse cell types for the neocortex. An initial period of excitatory neuron production is followed by a later phase in which the progenitor pool switches to making oligodendrocytes, astrocytes and olfactory bulb interneurons. Several studies have implicated Shh signaling in this transition, but the molecular mechanisms downstream of Shh are not fully understood. In this study, we analyzed scRNA-seq datasets to uncover transcriptional changes as progenitors transition to making OPCs in response to Shh. We found that 1) Shh signaling promotes a pre-OPC transcriptional state that emerges in progenitors prior to OPC appearance; 2) this pre-OPC state is characterized by enrichment of a gene regulatory network involving Ascl1 that has been shown to oppose an excitatory neuron transcriptional network; 3) the majority of cells generated by Ascl1^+^ progenitors in the late embryonic dorsal forebrain belong to the oligodendrocyte lineage. Together with previous work, these data lead us to propose a working model in which increased Shh signaling late in embryonic corticogenesis influences the balance between opposing transcriptional pathways to favor oligodendrogenesis over excitatory neuron fates (Fig. 11).

**Fig. 11.**
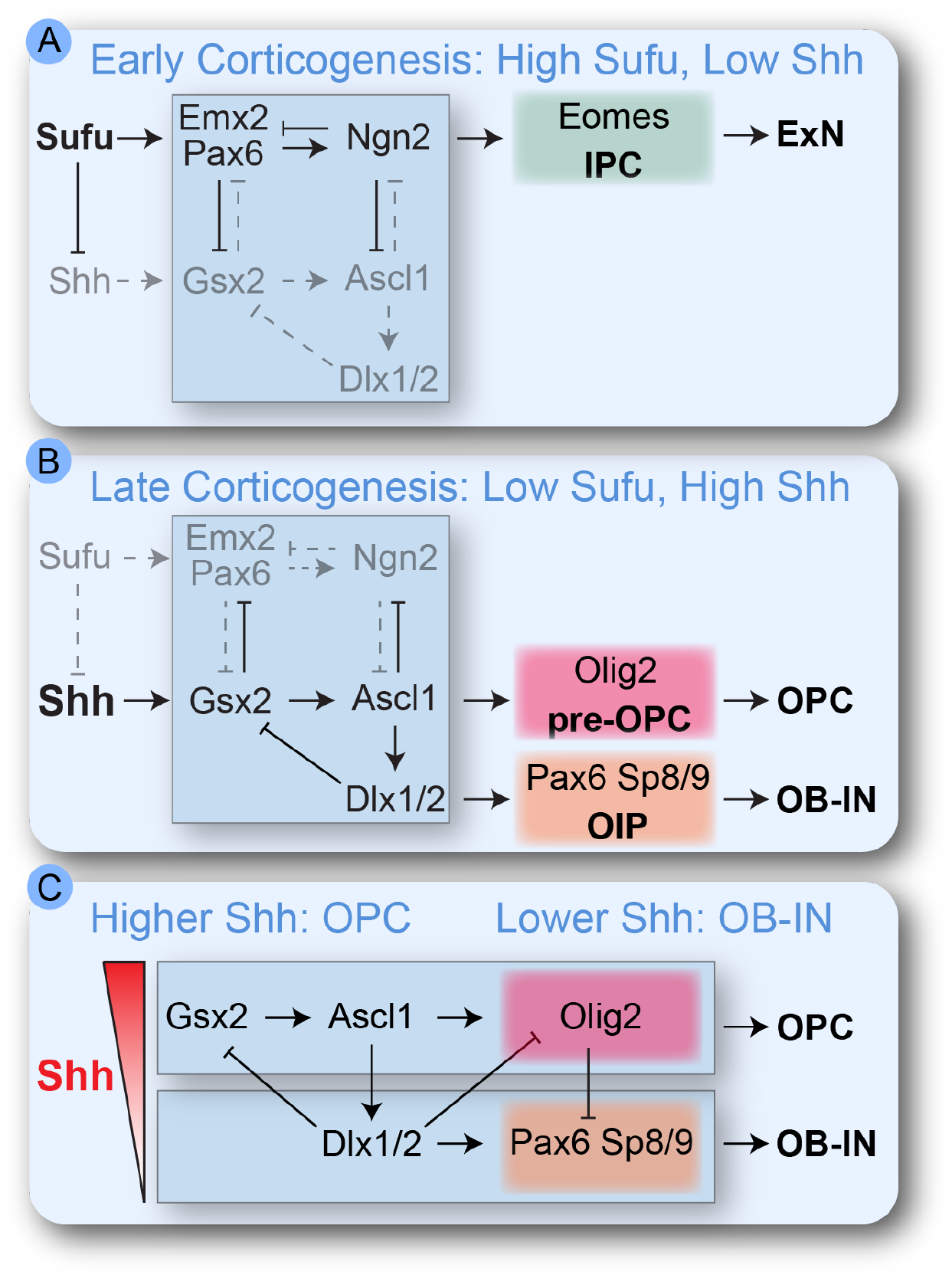
Working model for pre-OPC fate specification by Shh signaling. (A) Early in corticogenesis Shh signaling in the dorsal forebrain is low, due to low levels of Shh ligand and high expression of Sufu. This allows the *Emx2/Pax6* → *Neurog2* → Eomes pathway to repress ventral identity genes and drive specification and neurogenesis of excitatory neurons. (B) Later in corticogenesis Shh ligand increases in the dorsal forebrain while Sufu expression decreases. This promotes transcription of the ventral *Gsx2* → *Ascl1* → *Dlx1/2* & *Olig2* pathways, which repress dorsal excitatory neuron identity genes and drive specification of OPCs and OB-INs. (C) Levels of Shh signaling are also important for the fate decision between the OPC and OB-IN lineages, where the highest levels of Shh signaling promote oligodendrogenesis and lower Shh signaling promotes OB-IN generation through another cross-repressive transcriptional network.

### Multiple Temporal Roles for Shh Signaling in Cell Fate Specification

Throughout the CNS, Shh signaling is critical for setting up dorsal-ventral patterning of progenitor domains and for subsequently modulating diverse fates of their differentiated progeny, including oligodendrocytes (Andrews et al., 2019; Yabut and Pleasure, 2018). Recent studies have started to elucidate how levels and timing of Shh signaling are tightly regulated to control the identities of both progenitors and their progeny. Our lab uncovered a critical timing mechanism for Shh-mediated oligodendrogenesis in the developing neocortex (Winkler et al., 2018). Early in corticogenesis Shh signaling in the dorsal forebrain is low, due to weak expression of the ligand and high expression of the negative regulator, Suppressor of Fused (Sufu). This allows for proper dorsal-ventral patterning of forebrain progenitors. Later in corticogenesis, however, Shh signaling increases in the dorsal forebrain and is required for initiation of oligodendrogenesis. This is accomplished partly by decreased expression of Sufu (Yabut et al., 2015; Yabut et al., 2016), as well as increased Shh ligand brought in by interneurons migrating from the ventral forebrain (Winkler et al., 2018). In this way, tight control over the levels and timing of Shh signals confers both spatial patterning early in development and the later switch from neurogenesis to oligodendrogenesis.

Interestingly, recent studies together with our data presented here suggest that the molecular mechanisms involved in these dual roles might be very similar. Genetic ablation of the Shh repressor, Sufu, inthe early dorsal forebrain leads to overactivation of Shh-mediated transcription, resulting in patterning defects in dorsal progenitors (Yabut et al., 2015; Yabut et al., 2016; Yabut et al., 2020). The mis-specified dorsal progenitors ectopically express ventral identity genes, including *Gsx1/2, Dlxi/2, Olig2* and *Ascl1,* and downregulate dorsal identity genes like *Emx1/2* and *Pax6* (Yabut et al., 2015; Yabut et al., 2020). Thus, precocious activation of Shh signaling in dorsal progenitors activates a ventral transcriptional program. Our scRNA-seq analysis revealed a similar transcriptional network at later stages during oligodendrocyte lineage specification, in which *Gsx2*, *Dlx1/2* and *Ascl1* were all enriched in ShhPCs. Therefore, we hypotHes1ze that Shh signaling employs the same ventral transcriptional pathway in both early progenitor patterning and later OPC specification.

### Cross-repressive Gene Regulatory Networks for Neurogenesis and Oligodendrogenesis

A number of transcription factors work together in a network that confers dorsal identity and promotes specification of glutamatergic excitatory neurons for the neocortex. Emx2, Lhx2 and Pax6 are expressed in dorsal progenitors, where they are critical for establishing patterns of regional and areal identities and for maintaining progenitor proliferation (Hevner, 2006). Pax6 is also important for activating transcription of proneural genes *Neurog1* and *Neurog2* (Scardigli, 2003). Pax6 and Neurog1/2 are required for proper specification of a glutamatergic excitatory neuron identity (Schuurmans et al., 2004). Production of neocortical excitatory neurons most commonly proceeds through intermediate progenitor cells (Kowalczyk et al., 2009) that express the transcription factor Tbr2 (encoded by *Eomes*). Neurog2 regulates the transition from RGCs to IPCs by downregulating *Pax6* and initiating *Tbr2* expression (Kovach et al., 2013). Thus, a transcriptional pathway involving *Emx2/Pax6*→*Neurog1/2*→*Eomes* is central to the fate specification and genesis of glutamatergic excitatory neurons in the neocortex (Fig. 11A).

This dorsal transcriptional pathway participates in a cross-repressive network with an opposing pathway that confers ventral progenitor identities and GABAergic interneuron cell fates (Fig. 11A). In *Emx2*^/-^ and *Pax6^1^* mutant forebrains, dorsal progenitors ectopically express transcription factors in the ventral pathway, including *Gsx2, Ascl1, Dlx1* and *Dlx2,* and generate GABAergic inhibitory interneurons instead of glutamatergic excitatory neurons (Kroll and O’Leary, 2005). Emx2 and Pax6 can repress transcription of *Gsx2* (Cocas et al., 2011; Coutinho et al., 2011; Desmaris et al., 2018; Sun et al., 2015), and Pax6 and Neurog2 can similarly repress *Ascl1* expression (Gowan et al., 2001; Roybon et al., 2010; Sun et al., 2015). Conversely, knockout of *Gsx2* leads to ectopic expression of *Emx2*, *Pax6* and *Neurog2* in the ventral forebrain (Chapman et al., 2013; Corbin et al., 2000; Toresson et al., 2000; Yun et al., 2001), indicating that the ventral transcriptional network can also repress the dorsal pathway at multiple points.

Our analyses indicate that the ventral *Gsx2*→*Ascl1*→*Dlx1/2* transcriptional pathway is enriched in ShhPCs. This is consistent with some ShhPCs being committed to the OB-IN lineage (Zhang et al., 2020), as this pathway is important for generating OB-INs (Guo et al., 2019; Lindtner et al., 2019). In addition to its roles in neurogenesis, Ascl1 genetically interacts with *Olig2* and is critical for normal production of oligodendrocytes in the spinal cord and ventral forebrain (Nakatani et al., 2013; Parras et al., 2007; Sugimori et al., 2007). Olig2 can also repress *Pax6*, and both Pax6 and Tbr2 expression are reduced when Olig2 is overexpressed in the neocortex via electroporation (Lim et al., 2019). Based on these previous studies and the analyses presented here, we propose a model for how Shh signaling controls the switch from excitatory neurogenesis to production of OPCs and OB-INs (Fig. 11). Early in corticogenesis (Fig. 11A), Shh ligand expression is low dorsally, whereas Sufu expression is high in dorsal RGCs. This low Shh signaling state allows the *Emx2/Pax6*→*Neurog1/2*→*Eomes* pathway to repress ventral identity genes and drive specification and neurogenesis of excitatory neurons (Fig. 11A). Once the majority of excitatory neurons have been generated (Fig. 11B), Shh ligand increases in the dorsal forebrain while Sufu expression decreases, leading to increased Shh signaling in dorsal progenitors. High Shh signaling promotes transcription of the ventral *Gsx2*→*Ascl1*→*Dlx1/2*& *Olig2* pathways, which repress excitatory neuron identity genes and drive specification of OPCs and OB-INs (Fig. 11B).

A similar cross-repressive mechanism may be involved in the next fate decision between OPCs and OB-INs (Fig. 11C). The ShhPC cluster was further split into pre-OPCs and OB-IN precursors (OIPs). Pre-OPCs maintained higher levels of *Gsx2*, *Ascl1* and *Olig2* along pseudodevelopmental time, whereas OIPs increased their expression of *Dlx1/2, Pax6* and *Sp8/9.* Ascl1 can activate transcription of both *Olig2* (Vue et al., 2020) and *Dlx1/2* (Castro et al., 2011). In the ventral forebrain, Dlx1/2 negatively regulate Olig2-dependent OPC formation, whereas Ascl1 promotes OPC formation by restricting the number of Dlx1/2^+^ cells (Petryniak et al., 2007). In the developing chick forebrain, Dlx1/2 are necessary and sufficient to repress oligodendrocyte specification via transcriptional inhibition of *Olig2* (Jiang et al., 2020). Dlx1/2 also repress *Gsx2* (Guo et al., 2019) and activate *Sp8* and *Sp9* expression (Lindtner et al., 2019), which are essential for OB-IN maturation (Guo et al., 2019; Li et al., 2018). Pax6 is also required to generate a subset of OB-INs (Kohwi et al., 2005), and we found that *Pax6* levels are initially decreased in ShhPCs before increasing again specifically in the OB-IN lineage. Repression of *Pax6* by Olig2 (Jang and Goldman, 2011) might therefore be important for maintaining an OPC identity over that of OB-INs. Olig2 also activates *Olig1* expression and Olig1 directly represses *Dlx1/2* to inhibit interneuron production (Silbereis et al., 2014). Together, these studies raise the possibility that the relative levels of these cross-repressive transcription factors determine final fate commitment to either OPC or OB-IN lineages (Fig. 11C). Interestingly, we found that transcripts involved in Shh signaling were higher in the OPC lineage compared to the OB-IN lineage, suggesting that the levels of Shh may again be important for determining this binary fate choice. We propose that ShhPCs experiencing the highest levels of Shh signaling commit to the OPC lineage, whereas those with lower Shh signaling are fated to become OB-INs (Fig. 11C).

### Role of Ascl1 in Specifying Oligodendrocyte Fate Downstream of Shh

Our analyses revealed *Ascl1* as a marker of ShhPCs, consistent with experimental data that Shh signaling promotes Ascl1 expression (Voronova et al., 2011; Wang et al., 2007). Although Ascl1 is most highly expressed in ventral forebrain progenitors, we and others have shown that Ascl1 is expressed at low levels in dorsal progenitors (Britz et al., 2006; Fode et al., 2000; Winkler et al., 2018). Ascl1 is an early marker of the oligodendrocyte lineage in other CNS regions (Nakatani et al., 2013; Vue et al., 2014), and we confirm that Ascl1 is expressed in dorsal RGCs prior to expression of Olig2. We also previously demonstrated that dorsal Ascl1 expression requires Shh signaling (Winkler et al., 2018). Together, these data indicate that *Ascl1* is one of the first genes upregulated in the oligodendrocyte lineage in response to Shh. Previous lineage-tracing studies indicated that neocortical cells derived from the Ascl1 lineage were oligodendrocytes and ventrally-derived interneurons, but not astrocytes or excitatory neurons (Kim et al., 2008; Kim et al., 2011). Here, we combined directional *in utero* electroporation with short-term *Ascl1-CreERT_2_* lineage tracing to show that the majority of cells in the dorsal Ascl1 lineage were OPCs. The identities of the ~30% that are not OPCs remain to be determined, but transcriptomics and previous lineage analyses indicate that many are likely OB-IN neuroblasts.

Interestingly, a recent preprint found that E12.5 dorsal progenitors fall into one of four categories based on their expression of Ascl1 and Neurog2 (Han et al., 2020). The majority of progenitors were double-negative or Neurog2^+^, but a small proportion expressed Ascl1 or Ascl1 and Neurog2 together. They found by scRNA-seq that the Ascl1^+^ cells were enriched for oligodendrocyte and interneuron genes and were biased towards generating OPCs when cultured *in vitro*. Their *in silico* modeling indicated that “knocking out” Ascl1 perturbed a major gene regulatory network in Ascl1^+^ cells. These results are compatible with our *in vivo* lineage tracing and further support a critical role for Ascl1 in specifying the oligodendrocyte lineage in the dorsal forebrain.

Another interesting finding from Han *et al.* is that the Ascl1;Neurog2 double-negative and the Ascl1^+^ progenitor subsets were the most RGC-like, whereas the Neurog2^+^ and double-positive cells expressed higher levels of IPC and neuronal genes. We observed a similar pattern in our analyses, in which Ascl1^+^ShhPCs expressed several RGC markers and exhibited weaker RNA velocities compared to more differentiated cells. This connection between pre-OPCs and RGCs was even stronger in the human dataset, in which the pre-OPC cluster shared several markers with outer radial glia (oRG). Indeed, a recent study showed that oRGs are lineally related to pre-OPCs in the human neocortex (Huang et al., 2020). It will be interesting to test in the future whether this relationship between pre-OPCs and RGCs has any functional connection to the long-term proliferative capacity of OPCs, and whether Shh signaling is involved.

## Conclusion

We propose a model in which Shh signaling promotes an oligodendrocyte fate by influencing the balance between ventral and dorsal identities in a cross-repressive gene regulatory network. Future studies should shed light on the specific functions of individual transcription factors in this process, in particular the roles of Ascl1. It will also be interesting to determine whether a similar mechanism is at work during OPC production in other contexts, for example in the developing spinal cord or in the postnatal brain. Further investigations into oligodendrocyte development in the embryonic brain may also reveal insights into OPC responses to disease or injury in the postnatal brain, and potential strategies for treatment.

## MATERIALS AND METHODS

### Computational Analysis

#### Software

For all analyses we used a combination of Python (version 3.7.4) and R (version 3.6.2) packages and integrated R into Jupyter Notebook using the rpy2 interface (version 3.0.5) and anndata2ri (version 1.0.4) package (https://github.com/theislab/anndata2ri), which automatically converts from an AnnData object to a SingleCellExperiment object.

#### Samples

##### Shh-IUE Mouse Cortex

FASTQ files were downloaded from the National Center for Biotechnology Information BioProjects Gene Expression Omnibus (GEO: GSE140817; Shh-IUE samples) (Zhang et al., 2020) and processed using Cell Ranger (version 2.1.1) with mm10 genome assembly to generate unique molecular identifier (UMI) gene count matrices. An aggregated matrix was also generated by downsampling of the mapped reads in each sample to the same depth as the sample with the lowest read count using cellranger aggr. The aggregated matrix was processed using Scanpy (version 1.4.6) (Wolf et al., 2018) to perform initial quality control filtering, normalization, and clustering.

##### WT Mouse Neocortex

For embryonic ages E11.5, E13.5, and E15.5 we downloaded provided tab-delimited text files containing raw digital gene expression for all cells (GEO: GSE107122) (Yuzwa et al., 2017). For embryonic ages E12.5, E16.5, and E18.0 we downloaded provided developmental data as a single file (dev_all.loom) from the Mouse Brain Atlas website compiled by the Linnarsson lab (mousebrain.org). We then extracted the developmental timepoints of interest as separate h5ad files. Both the tab-delimited text files and h5ad files were read in and converted to AnnData objects for easy integration into our Scanpy pipeline for preprocessing and downstream analysis.

##### Human Neocortex

We downloaded the raw expression matrix for primary human cortex samples directly from the provided UCSC cell browser website (https://cells.ucsc.edu/?ds=organoidreportcard), as they were not yet available through the database of Genotypes and Phenotypes (dbGaP) at the time of analysis (GEO: GSE132672) (Bhaduri et al., 2020). We processed the raw expression matrix in R to selectively isolate samples from gestational week 18 (GW18). We used Seurat (version 3.1.5) to create a SeuratObject of the GW18 dataset, which we then converted to an AnnData object that we could easily integrate into our Scanpy pipeline for pre-processing and downstream analysis.

#### Pre-processing

Pre-processing and downstream analysis on the data sets was conducted following the steps outlined by Luecken and Theis (2019). For all data sets, genes were excluded if they were detectable in fewer than 5 cells. For <u>*Shh-IUE*</u> data, cells were removed if there were fewer than 1000 genes detected, a UMI count less than 2500, a UMI count more than 20,000, or if the proportion of UMIs mapped to mitochondrial genes was greater than 15%. We also excluded cells if the proportion of UMIs mapped to ribosomal genes was greater than 18% on a second pass through of the data, as we noticed that these cells represented stressed and/or dying cells. This resulted in 8154 cells (from 9080 total cells) after filtering. For <u>*Eii.5 WT*</u> data, cells were removed if there were fewer than 500 genes detected, a UMI count less than 700, a UMI count more than 10,000, or if the proportion of UMIs mapped to mitochondrial genes was greater than 5%, resulting in 1837 cells (from 2000 total cells) after filtering. For <u>*E13.5 WT*</u> data, cells were removed if there were fewer than 500 genes detected, a UMI count less than 500, a UMI count more than 10,000, or if the proportion of UMIs mapped to mitochondrial genes was greater than 20%, resulting in 1729 cells (from 2000 total cells) after filtering. For <u>*E15.5 WT*</u> data, cells were removed if there were fewer than 500 genes detected, a UMI count less than 750, a UMI count more than 20,000, or if the proportion of UMIs mapped to mitochondrial genes was greater than 15%, resulting in 4725 cells (from 5000 total cells) after filtering. For <u>*E12.5 WT*</u> data, cells were removed if there were fewer than 1000 genes detected, a UMI count less than 2000, a UMI count more than 10,000, or if the proportion of UMIs mapped to mitochondrial genes was greater than 2%, resulting in 2378 cells (from 2601 total cells) after filtering. For <u>*E16.5 WT*</u> data, cells were removed if there were fewer than 1000 genes detected, a UMI count less than 1000, a UMI count more than 15,000, or if the proportion of UMIs mapped to mitochondrial genes was greater than 20%, resulting in 2871 cells (from 3185 total cells) after filtering. For <u>*E18.0 WT*</u> data, cells were removed if there were fewer than 1000 genes detected, a UMI count less than 1000, a UMI count more than 10,000, or if the proportion of UMIs mapped to mitochondrial genes was greater than 20%, resulting in 4045 cells (from 4859 total cells) after filtering. For <u>*GW18*</u> data, cells were removed if there were fewer than 500 genes detected, a UMI count less than 1000, a UMI count more than 12,000, or if the proportion of UMIs mapped to mitochondrial genes was greater than 10%, resulting in 73623 cells (from 78453 total cells) after filtering.

Following filtering, the UMI counts for all data sets were normalized via scran normalization (version 1.14.6) and log-transformed with an offset of 1. For the <u>*GW18*</u> data, following normalization we used ComBat batch correction (Johnson et al., 2007), as cells were collected from various regions across the neocortex and were clustering based on region. To reduce the dimensionality of each dataset the top 4000 highly variable genes were extracted for further processing. Dimensions of each dataset were further reduced by principal component analysis (using the top 50 principal components) and visualized using Uniform Manifold Approximation and Projection (UMAP). Finally, we used a gene list from Macosko et al. (Macosko et al., 2015) to score the cell cycle effect in each data set and classify cells by cell cycle phase.

#### Cluster identification and annotation

Graph-based clustering was performed on the highly variable gene data for each data set, dimensionality reduced by Principal Component Analysis (PCA), and embedded into K-Nearest Neighbors (KNN) graph using the Louvain algorithm with a resolution of 1.0. Clusters were annotated using a combination of marker genes generated by differential expression testing (using the Wilcoxon rank-sum test) and by visualizing expression of known marker genes of various cortical cell types (see Table S1 for mouse genes, Table S8 for human genes). We analyzed the expression of both individual genes as well as marker gene sets that we compiled based on differential expression testing and published literature (Tables S1 and S8). After cluster annotations, we repeated the clustering as described on a subset of cells that included all progenitor cell types (and immature olfactory bulb interneurons (OB-IN) in the <u>*Shh-IUE*</u> data set). Here we regressed out cell cycle effects to avoid progenitors clustering based on cell-cycle stage. For the <u>*Shh-IUE*</u> data, in order to further subcluster the Shh-responding progenitor cell (ShhPC) cluster we performed Louvain clustering on just the ShhPC cluster using a resolution of 1.0.

#### Functional enrichment analysis

We used the g:Profiler web server (https://biit.cs.ut.ee/gprofiler/gost) to perform functional enrichment analysis on the ShhPC1 and ShhPC3 clusters. Lists of differentially expressed genes in ShhPC1 versus rest (Table S4) and ShhPC2 versus rest (Table S6) were generated using built-in features of Scanpy (using the Wilcoxon rank-sum test). We used g:Profiler to map these gene lists to known functional information sources and detect statistically significantly enriched terms (Tables S5 and S7).

#### Trajectory analysis

The following downstream analysis was only performed in the <u>*Shh-IUE*</u> data set. Cell lineage trajectory within the progenitor subset was analyzed using Slingshot (version 1.5.1) (Street et al., 2018). Dimensionality was reduced by PCA, and we used an unbiased approach for lineage construction by only specifying a start cluster (radial glial cells, or RGCs, the most stem-like progenitor cluster). This predicted two distinct lineage trajectories (an oligodendrocyte lineage and OB-IN lineage) which we visualized using Slingshot visualization tools.

We also did RNA velocity analysis as a second, independent method of trajectory analysis. We used the Python implementation of velocyto (version 0.17.13) (La Manno et al., 2018), using the basic ‘run’ subcommand with our Cell Ranger output and mm10 genome assembly. We also opted to mask expressed repetitive elements using the mm10 expressed repeat annotation file downloaded from the UCSC genome browser. We analyzed the output loom file using scVelo (version 0.2.2) (Bergen et al., 2020), and merged the loom file with our Scanpy-processed and -annotated progenitor subset. The resulting abundance of spliced to unspliced RNA was 0.73:0.27. We set the minimum number of counts (both unspliced and spliced) required for a gene to 1 and filtered out 1446 genes. We used default arguments in order to compute moments for velocity estimation.

#### Trajectory-based differential expression analysis

We used tradeSeq (version 1.1.16) (Van den Berge et al., 2020) to infer differential gene expression between the lineages predicted by our Slingshot analysis. We used k=8 knots to fit the tradeSeq negative binomial generalized additive model for each gene in the Slingshot dataset and estimated one smoother for each of our two lineages (default parameters). We used built-in tools provided by tradeSeq to generate lists of genes that were differentially expressed between the oligodendrocyte and OB-IN lineages between knots 2 and 5 (Table S3), as this is when the lineages first began to bifurcate. Smoothers were visualized using built-in tools.

#### Experimental Methods

##### Mice

The following mice were obtained from The Jackson Laboratory: B6 (*C57BL/6J;* stock no. 000664); *Ai9 (B6.Cg-Gt(ROSA)26Sor^tm9(CAG^-^tdT¤mat¤)Hze^/J);* stock no. 007909); *Ascl1-CreERT2 (Ascl1^tmι.1(Cre/ERT2)ĩei¤^/J);* stock no. 012882); *Emx1-Cre (B6.i29S2-Emx1^tmι(cre)Krj^/J,* stock no. 005628); *R26-NZG* [FVB.Cg*-Gt(ROSA)26Sor^tmι(CAG-lacZ,-EGFP)Glh^/J,* stock no. 012429]. *Emx1-^Cre/Cre^* homozygous mice were crossed with *R26-^NZG/NZG^* homozygous mice to generate double heterozygous embryos for analysis. *Ascl1^+/CreERT2^* heterozygous males were crossed to *Ai9^fl/*fl*^*homozygous females to generate pregnant dams for electroporation and tamoxifen injection. Animals were maintained according to the guidelines from the Institutional Animal Care and Use Committee of the University of Colorado – Anschutz Medical Campus. Sex of experimental embryos was not determined in our experiments.

##### Tissue preparation

Embryonic brains were fixed in 4% paraformaldehyde (PFA) for 1 h at room temperature (RT). Postnatal day 0 (P0) brains were fixed in 4% PFA overnight at 4 degrees C. For immunohistochemistry, brains were sectioned coronally at 50–100 μm with a vibrating microtome. For RNAscope® (ACD) mRNA in situ hybridization, brains were cryoprotected in 30% sucrose at 4 degrees C overnight and sectioned on a cryostat at 12 μm.

##### Immunohistochemistry

Free-floating vibratome sections were blocked with 10% donkey serum and 0.2% Triton-X in 1X PBS for 2 h at RT. After 2 h, the blocking solution was removed and sections were incubated with primary antibodies in 10% donkey serum in 1X PBS overnight at 4 degrees C, and then washed at RT with 1X PBS three times for 5 min each. After washing, sections were incubated with secondary antibodies in 10% donkey serum in 1X PBS for 1 h at RT, and then washed again with 1X PBS three times for 5 min each. Sections were mounted on slides with ProLong Diamond Antifade Mountant (Invitrogen P36961). Images were captured using a LSM900 Zeiss laserscanning confocal microscope in Airyscan 2 Multiplex 4Y mode at 20x (PlanApo 20x/0.8 M27, Zeiss 420650-9902-000) or 40x (C-Apo 40x/1.20 W Corr M27, Zeiss 421767-9971-000) magnification. Antibodies used for immunostaining wild-type B6 brains were: rabbit anti-Ascl1 (1:1000; Abcam [EPR19840] ab211327), mouse anti-Olig2 (1:500; Millipore MABN50; RRID:AB_10807410), chicken anti-β-gal (1:2000; Abcam ab9361; RRID:AB_307210). Rabbit anti-Olig2 (1:500; Millipore AB9610; RRID:AB_570666) was used for immunostaining electroporated brain sections. Donkey secondary antibodies conjugated to Alexa Fluor 488, Rhodamine Red-X, or Alexa Fluor 647 were purchased from Jackson ImmunoResearch and used at 1:500.

##### RNAscope® mRNA *in situ* hybridization

mRNA in situ hybridization was performed using the RNAscope® Multiplex Fluorescent Reagent Kit v2 (Advanced Cell Diagnostics) with an *Ascl1* probe (RNAscope® Probe – Mm-*Ascl1-*C2; Advanced Cell Diagnostics, catalog no. 313291-C2) and an *Emx1* probe (RNAscope® Probe – Mm-*Emx1*; Advanced Cell Diagnostics, catalog no. 312261). Signals were visualized with Opal 520 (Akoyo Biosciences FP1487001KT) and Opal 570 (Akoyo Biosciences FP1488001KT). The assay was performed in accordance with the manufacturer’s instructions, except that slides were steamed in target retrieval buffer lying flat horizontally instead of vertically, to prevent sections from sliding. Sections were counterstained with DAPI to label nuclei.

##### *In Utero* Electroporation and Tamoxifen Injection

A *piggyBac* (PB) transposase system was used for *in utero* electroporation experiments to permit stable integration of the reporter plasmid into electroporated progenitors (Silbereis et al., 2014). pPB-nuc.mT-Sapphirie was generated by removing the Lox-STOP-Lox cassette from pPB-STOP-nuc.mT-Sapphire (García-Moreno et al., 2014) by incubating 0.25 μl of plasmid with 1 μl of recombinant Cre recombinase (New England Biolabs M0298S) according to manufacturer’s instructions. *In utero* electroporations were performed as described (Winkler et al., 2018). Briefly, timed pregnant mice (E15.5) were anesthetized and their uterine horns exposed. 1 μl of endotoxin-free plasmid DNA was injected into the embryos’ lateral ventricles at the following concentrations: pPB-nuc.mT-Sapphire 1 mg/mL, CMV-mPB 0.3 mg/mL (Winkler et al., 2018). 5 electrical pulses separated by 950 ms were applied at 45 V. Tamoxifen injections of the pregnant dams were performed immediately after the electroporated embryos were returned to the abdominal cavity and the dams were sutured. Tamoxifen (Sigma-Aldrich T5648) was dissolved in corn oil (Sigma-Aldrich C8267) at (10 mg/ml) and administered by intraperitoneal injection of 2 mg. Embryos were allowed to develop *in utero* for the indicated time. For quantification of electroporated cells that were Olig2^+^, all nuc.mT-Sapphire;tdTomato double-positive electroporated cells in each entire section were analyzed for Olig2 expression by IHC with rabbit anti-Olig2.

## Acknowledgements

We thank members of the Pediatrics Department Section of Developmental Biology and the RNA Biosciences Informatics Fellows for helpful discussions.

## Author contributions

Conceptualization: C.C.W., S.J.F.; Methodology: C.C.W., L.N.T., E.P.M., F.G.-M., S.J.F.; Formal analysis: C.C.W.; Investigation: C.C.W., L.N.T., E.P.M., S.J.F.; Resources: F.G.-M.; Data curation: C.C.W.; Writing - original draft preparation: C.C.W., S.J.F.; Writing - review and editing: L.N.T.; Visualization: C.C.W., L.N.T., S.J.F.; Supervision: C.C.W., S.J.F.; Project administration: C.C.W., S.J.F.; Funding acquisition: C.C.W., S.J.F.

## Data availability

Jupyter Notebook workflows available upon request.

